# Transmission, tropism and biological impacts of torix *Rickettsia* in the common bed bug *Cimex lectularius* (Hemiptera: Cimicidae)

**DOI:** 10.1101/2020.09.20.305367

**Authors:** Panupong Thongprem, Sophie EF Evison, Gregory DD Hurst, Oliver Otti

## Abstract

The torix group of *Rickettsia* have been recorded from a wide assemblage of invertebrates, but details of transmission and biological impacts on the host have rarely been established. The common bed bug *(Cimex lectularius)* is a hemipteran insect which lives as an obligatory hematophagous pest of humans and is host to a primary *Wolbachia* symbiont and two facultative symbionts, a BEV-like symbiont, and a torix group *Rickettsia*. In this study, we first note the presence of a single *Rickettsia* strain in multiple laboratory bed bug isolates derived from Europe and Africa. Importantly, we discovered that the *Rickettsia* has segregated in two laboratory strains, providing infected and uninfected isogenic lines for this study. Crosses with these lines established transmission was purely maternal, in contrast to previous studies of torix infections in planthoppers where paternal infection status was also important. Fluorescence *in-situ* hybridization analysis indicates *Rickettsia* infected in oocytes and bacteriomes, and other somatic tissues. There was no evidence that *Rickettsia* infection was associated with sex ratio distortion activity, but *Rickettsia* infected individuals developed from first instar to adult more slowly. The impact of *Rickettsia* on fecundity and fertility were investigated. *Rickettsia* infected females produced fewer fertile eggs, but there was no evidence for cytoplasmic incompatibility. These data imply the existence of an unknown benefit to *C. lectularius* carrying *Rickettsia* that awaits further research.

## INTRODUCTION

Symbioses between insects and bacteria are both common and important. Bacterial symbionts impact on the biology of their host individual, and also by extension affect the ecology and evolution of their host (1, 2). Effects are diverse, with beneficial effects ranging from nutritional (e.g. anabolism or digestion) and protection against multiple different forms of natural enemies and even xenobiotics (3-5). Parasitic interactions are also known, commonly associated with sex ratio distortion activity selected for as a consequence of the maternal inheritance of the symbiont (6).

The genus Rickettsia (alpha-proteobacteria), has emerged as an important associated of insects. Members of this genus were classically considered as causative agent of arthropod-borne rickettsioses, which threaten livestock and human health (7, 8). These zoonotic pathogens are endosymbionts of ticks, mites, fleas and lice (9). Following a bite, the bacteria disseminate into the blood of mammals where it causes disease such as typhus and spotted fever (7, 9). In 1994, however, it was first discovered in ladybirds (*Adalia bipunctata*) that *Rickettsia* can exist strictly as vertically transmitted endosymbionts of arthropods (10), with no mammalian transmission. These *Rickettsia* symbionts are known to present a significant selection pressure on their insect hosts by, for example, altering reproductive success and distorting sex ratio by inducing male-killing in ladybirds (10) and parthenogenesis in a parasitoid wasp (11). Unlike *Cardinium, Rickettsiella* and *Wolbachia, Rickettsia* has never been involved in causing cytoplasmic incompatibility in arthropods (12-14). With regard to host fitness, some *Rickettsia* strains may upregulate host immunity and thereby improve the host defence against pathogens (4).

In 2002, a new group of Rickettsia were discovered during research on *Torix tagoi* leeches. A rickettsia was found that was associated with larger leech size (15, 16). The *Rickettsia* was a sister group to all other Rickettsia described previously, and the clade were named ‘torix *Rickettsia’* (15). Torix *Rickettsia* have since been found across multiple arthropod taxa and seem to be widespread and especially common in species associated with freshwater, e.g. *Culicoides* midges (17), dytiscid water beetles (18), Odonata (19) and Amphipoda (20), but has also been detected in some terrestrial arthropods, e.g., Araneidae (21), Siphonaptera (22) and Hemiptera (23). Whilst we now understand symbioses between invertebrates and torix Rickettsia are common, much less is known of their biological significance. This knowledge deficit arises largely from the lack of a good laboratory model systems in which inheritance and biological impact can be measured.

In this paper, we characterize the patterns of inheritance and biological impact of torix *Rickettsia* in the common bed bug (*Cimex lectularius*). This species is in the order Hemiptera, belonging to the family Cimicidae, all members of which are ectoparasites of warm-blooded animals (24). *C. lectularius* is a human parasite and its global pest status has medical, social and economic impacts (25, 26). Being an obligate haematophage, *C. lectularius* have evolved a special organ called a ‘bacteriome’ that harbours *Wolbachia*, which live as a primary endosymbiont and synthesise B-vitamins to supplement the host’s deficient diet (3). In some individuals, *Wolbachia* is found alongside a facultative gamma-proteobacterium, also known as BEV-like symbiont (3, 27). The impact of this symbiont is not currently understood.

A recent PCR-based screen by Potts et al. (28) revealed *Rickettsia* associated with natural populations of *C. lectularius* in both the UK and the USA. Partial citrate synthase gene (*gltA*) sequences showed that this strain is closely related to the *Rickettsia* found in the flea *Nosopsyllus laeviceps* (22). Recent work by ourselves has indicated *C. lectularius* genomic DNA, samples from Duron et al. in 2008 (29), also carries a *Rickettsia* symbiont. The analysis revealed the presence of torix *Rickettsia* in multiple individuals from one laboratory strain, F4.

To explore the system of torix *Rickettsia* association with *C. lectularius*, we undertook PCR assays to investigate the genetic diversity and prevalence of these *Rickettsia* strains in *C. lectularius* cultures originally collected from various locations across the UK, Europe and Africa. We isolated two laboratory lines where *Rickettsia* infection had segregated, and used these to analyse the transmission, tissue tropism and biological impacts of *Rickettsia* infection. These results indicate a maternally inherited symbiont with a broad somatic/germline distribution, that does not impact host sex ratio or generate cytoplasmic incompatibility. Impacts on host development and reproduction are minor, indicating there is an as yet unelucidated impact on the host.

## MATERIALS AND METHODS

### Prevalence of torix *Rickettsia* in *C. lectularius* populations and cimicid allies

One male and one female adult bed bug from each of 21 lab populations maintained at the University of Bayreuth (Table 1) were sent in 2.0 ml absolute ethanol tubes (one pair/tube) to the lab at the University of Liverpool for DNA extraction. These populations were collected from different areas of Europe and Africa in different years (Table 1). Two populations are of unknown origin in the wild. One has been maintained at the Universities of Bayreuth and Sheffield for >20 years and before that for >40 years at the London School of Hygiene and Tropical Medicine. The other population was received from Bayer (Germany) in 2006.

**Table 1:** *Rickettsia* infection in *Cimex lectularius* stock populations in Bayreuth lab. Males in the brackets (M) indicate the infection status were detected later when these populations were screened with more individuals. Asterisks after a population name indicate that these populations were selected for transmission mode experiment and established as the isofemale lines.

| Population name | Origin of place | Year established<br>in Sheffield | Male /Female (M/F)<br>positive |
| --- | --- | --- | --- |
| **H1** | Budapest, Hungary | 2010 | neither |
| **C1** | Coventry, UK | 2010 | neither |
| **Bayer** | Germany | 2006 | neither |
| **S1\*** | London School of Tropical Medicine | 1996 | (M)/F |
| **London heavy** | London, UK | 2006 | neither |
| **F11** | London, UK | 2006 | neither |
| **F4\*** | London, UK | 2006 | M/F |
| **F10** | London, UK | 2006 | neither |
| **YMCA** | London, UK | 2010 | neither |
| **K12** | Nairobi, Kenya | 2008 | M/F |
| **K4** | Nairobi, Kenya | 2008 | M/F |
| **K22** | Nairobi, Kenya | 2010 | M/F |
| **K25** | Nairobi, Kenya | 2010 | M/F |
| **K26** | Nairobi, Kenya | 2010 | M/F |
| **K19** | Nairobi, Kenya | 2010 | M/F |
| **K20** | Nairobi, Kenya | 2010 | M/F |
| **K7** | Nairobi, Kenya | 2008 | M/F |
| **K5** | Nairobi, Kenya | 2008 | M/F |
| **K30** | Nairobi, Kenya | 2010 | M/F |
| **BG1** | Sofia, Bulgaria | 2010 | (M)/F |
| **K17** | Watamu, Kenya | 2010 | neither |

The samples were rinsed with absolute ethanol and left at room temperature until the samples dried. To avoid contamination with gut microbes, bed bugs were decapitated with sterilised forceps and only the head and/or the upper part (from the head to the thorax, including legs) taken for DNA extraction. Genomic DNA was extracted from the selected body part using Promega Wizard® Genomic DNA Purification kit (A1120, Promega, UK) and DNA dissolved with 100 ul of molecular water and stored in −20°C for the future use.

In addition, DNA template was obtained from various species of cimicids from the recently published bed bug phylogeny by Roth et al. (2019) (Table 2). For each species, we received 10 µl extracted genomic DNA in 0.2 µl tubes from Steffen Roth (University Museum of Bergen, Norway) and Klaus Reinhardt (TU Dresden, Germany), which we have stored in −20°C until used. Voucher specimens of some species are stored in the collection of the University Museum of Bergen (ZMNB), Norway.

**Table 2.** *Rickettsia* infection in cimicid allies. The infection found in all individuals of *Afrocimex constrictus* but absent in other tested species.

| Subfamily | Species | N | Locality | Year of collection | Infection |
| --- | --- | --- | --- | --- | --- |
| **Afrocmimicinae** | ***Afrocmimex constrictus*** | **3** | **Kenya** | **2005** | **+** |
| Primicinae | *Bucimex chilensis* | 1 | Chile | 2013 | - |
|  | *Primicimex cavernis* | 1 | Mexico | 2015 | - |
| Haematosiphoninae | *Ornithocoris pallidus* | 1 | USA, South Carolina | 2010 | - |
|  | *Acanthocrios furnarii* | 1 | Brazil | 2010 | - |
|  | *Psitticimex uritui* | 1 | Argentina | 2008 | - |
|  | *Cyanolicimex patagonicus* | 1 | Argentina | 2003 | - |
|  | *Cimexopsis nyctalis* | 1 | USA | 2016 | - |
|  | *Hesperocimex sonorensis* | 1 | Mexico | 2017 | - |
|  | *Hesperocimex coloradensis* | 1 | Los Alamos County, N.Mex. | 1971 | - |
| Cimicinae | *Paracimex inflatus* | 1 | Kavieng, Papua New Guinea | 1996 | - |
|  | *Paracimex borneensis* | 1 | Borneo | 2015 | - |
|  | *Cimex pipistrelli* | 1 | Hanau, Germany | 2004 | - |
|  | *Cimex hemipterus* | 1 | Taiwan | Before 1966 | - |
|  | *Cimex hirundinis* | 1 | Switzerland | N/A | - |
| Cacodminae | *Aphrania elongata* | 1 | Senegal | 2012 | - |
|  | *Aphrania vishnou* | 1 | Phnom Penh, Cambodia | 1952 | - |
|  | *Cacodmus villosus* | 1 | Kenya | 2005 | - |
|  | *Loxapsis malayensis* | 1 | Tasik Bera, Pahang, Malaysia | 1962 | - |
|  | *Leptocimex duplicatus* | 1 | Israel | 2002 | - |
|  | *Leptocimex boueti* | 1 | N/A | N/A | - |

Initially, all DNA samples were checked for their quality using the invertebrate mtDNA barcoding primers C1J_1718 (30) and HCO_2198 (31) (Supplementary Table 1) that amplified a fragment of approximately 380 bp of the cytochrome oxidase subunit I (*COI)* gene of *C. lectularius*. For the samples that passed quality control, we assessed the presence of *Rickettsia* infections using two *Rickettsia*-specific primer pairs, targeting 16S rRNA and citrate synthase gene *gltA* (16SrRNA: Ri170_F and Ri1500_R (18); *gltA*, RiGltA405_F and RiGltA1193_R (17), Supplementary Table 1). These primer pairs for *Rickettsia* have been tested to specify known *Rickettsia* groups which do not cross amplify other Rickettsiales (17-19). If we found only one sex to be *Rickettsia*-positive in a *C. lectularius* population in the initial screen, we screened more individuals (4-10 samples mixed males and females) to verify the infection status in the other sex.

### Relatedness of strains

The *16S rRNA* and *gltA* amplicons from PCR assays were cleaned with the ExoSAP-IT kit (E1050, New England Biolabs, US) and Sanger sequenced. The sequence chromatograms were trimmed and edited in UGENE (32). All the sequences were exported to fasta format and searched against other *Rickettsia* strains on NCBI database to find close relatives ascertained by BLAST homology. The sequence of these markers from closely related strains from other invertebrate hosts were retrieved, and the relatedness within the torix group estimated. Other *Rickettsia* strains from other clades, e.g., *Rickettsia bellii* and vertebrate pathogens were selected to represent the sister group to the torix clade. *Occidentia massiliensis* was use as the outgroup for both topologies. All the selected sequences were aligned with *Rickettsia* sequence of *C. lectularius* using MUSCLE algorithm with its default setting in MEGA X (33, 34). The ML phylogeny for the both genes were estimated in MEGA X with 1000 rapid bootstrap replicates under T92+I and K2+I model for *gltA* and 16S gene respectively.

### Transmission mode experiment

To investigate the vertical transmission mode of torix *Rickettsia*, we used two bed bug lab populations, S1 and F4, in which we found the sporadic infections with infected and uninfected individuals throughout the populations. We randomly selected males and females to establish 65 and 49 mating pairs for S1 and F4 respectively, from which we reared offspring.

The parents and 5-10 randomly selected first instar nymphs were screened for torix infection status using the PCR assays as described above. Nymphs were tested individually to gain insight into vertical transmission efficiency, and whole bodies were used for template. Then we assessed the impact of parental infection status (mother infected, father infected) on progeny infection status.

### Isofemale lines and bed bug culture

Based on the infection status of offspring from the transmission mode experiment, we established four *Rickettsia*-free (R−) and four *Rickettsia*-infected (R+) isolines for each of the F4 and S1 populations. These isofemale lines of known *Rickettsia* infection state were then kept under constant conditions, in an CT room at 26±1°C, at about 70% relative humidity with a cycle of 12L:12D. We additionally tested all isofemale lines for the BEV-like bacterium infection using a PCR assay as described in Degnan et al (35) (Supplementary Table 1). New generations were set up regularly, i.e. at a 6- to 8-week interval. Each new generation was started with randomly picked virgin female and virgin male. All bed bugs were maintained in the CT room with the conditions described as above. All individuals in our study were virgin prior to experiments. The feeding, maintenance and generation-of-virgin-individuals protocols follow Reinhardt et al. (36)

### Fluorescence *in situ* hybridization (FISH)

To localise the torix *Rickettsia* and other symbionts within the *C. lectularius* body, we used the FISH technique adapted from Sakurai et al. (37). We investigated the bacteriome and reproductive tissue in virgin male and female adults, as well as the whole body of first instar nymphs from the torix-free and torix-infected F4 and S1 lines. Tissues were dissected in 0.5M PBS at pH 7.4 and preserved immediately in Carnoy’s solution (chloroform: ethanol: glacial acetic acid = 6:3:1) overnight. The nymphs were preserved in the solution without dissection. All tissue samples were cleared by incubating in 6% H_2_O_2_ in ethanol for 12 hr, for the whole-body nymph was incubated at least 24 hr or until the body was transparent. We then used a tungsten micro-needle to make micropores in the nymph cuticle to allow the fluorescence probes to pass through the cuticle during the hybridization step. The samples were hybridised by incubating the tissues overnight in a hybridization buffer (20mM Tris-HCl pH 8.0, 0.9M NaCl, 0.01% Sodium dodecyl sulfate 30% formamide) with 10 pmol/ml of these symbionts rRNA specific probes *Rickettsia* (38), *Wolbachia* (3) and Gamma proteobacteria (BEV-like symbiont) (3) (see the probe sequences in Supplementary Table 1). We also used nuclei fluorescence staining, Hoechst 33342 (H1399, Invitrogen, Carlsbad, USA), to visualise the bed bug tissues. After incubation, tissues were washed in buffer (0.3M NaCl, 0.03 M sodium citrate, 0.01% sodium dodecyl sulfate) and mounted onto a slide using VECTASHIELD® Antifade (H-1000, Vectorlabs, UK) as a mounting medium. Slides were then observed under a confocal microscope, 880 Bio AFM (on 880 LSM platform, ZEISS, Germany).

### Development time and sex ratio produced by *Rickettsia* infected and uninfected individuals in a common garden experiment

A 7-day-old virgin male and female from each isofemale line were put together in a pot and allowed to mate. Once offspring hatched from the eggs laid, we collected ten 1^st^ instar nymphs from each pot (N = 160). We then prepared eight fresh pots with a filter-paper and randomly assigned each group of ten *Rickettsia*-infected and ten *Rickettsia*-free nymphs to a pot and presented them with the opportunity to feed every three days. As soon as the first 5^th^ instar nymph was observed in a pot, eclosion was checked every day. Freshly eclosed adults were then removed from the pot and post hoc screened for Rickettsia infection status using the PCR method described above. The number of days between placement into the pot and the last hatching event, i.e. removal from the pot, represents the development time. The sexes of individuals were determined when the bugs reached the adult stage. Sex ratio (number of female:male) was calculated and compared between the two infection status, *Rickettsia*-infected and *Rickettsia*-free individuals, which were identified with PCR assays as described above.

### Effect of *Rickettsia* infection on fecundity and cytoplasmic incompatibility

To measure the effect of *Rickettsia* infection on fecundity we used a full factorial crossing scheme of female x male i.e. R+ x R+, R+ x R−, R− x R+ and R− x R−. For this, we prepared same-aged individuals by putting a 7-day-old virgin male and female together in a pot and allowing them to mate and feed weekly. Every week we collected all the eggs and put them in a fresh pot, which was fed weekly until 5^th^ instar nymphs were observed. Same-aged 5^th^ instars were then fed and placed into a 96-well plate until they reached adulthood. Seven-day-old virgin adults were then used in a full factorial crossing experiment. Prior to the experiment, females were fed twice, the last time on the day of mating, and males once, on the day of hatching. To avoid inbreeding effects, each isofemale line was crossed with every other line, but not with itself. To have equal sample sizes for within versus between *Rickettsia*-free and *Rickettsia*-infected crosses we randomly left one cross out. In this manner, we crossed each isofemale line with three *Rickettsia*-free lines and with three *Rickettsia*-infected lines (N = 96 crosses). Matings were staged, monitored and interrupted after 60s as described earlier in Reinhardt et al. (39). Interrupted matings were conducted to standardise sperm number because of the linear relationship between copulation duration and sperm number (40). A standardised sperm number was desirable since spermatozoa trigger the release of an oviposition-stimulating hormone from the *corpora allata* (41) and could potentially influence lifespan through differential egg production. The use of 60s standard matings also allows comparability with other studies.

After mating, the females were kept individually in 15ml plastic tubes equipped with a piece of filter paper for egg laying. Females were fed weekly and the number of fertile and infertile eggs counted in weekly intervals. Fertile and infertile eggs were distinguished, and the onset of laying and fertilization senescence determined following Reinhardt et al. in 2009 (42). The onset of infertility can be precisely obtained as the time point when the second infertile egg was laid, to allow for one accidental fertilization failure. The different of number of the fertile eggs were used to test the fecundity fitness.

To determine the presence of CI, we observed the number of fertile eggs produced by females from different crossing combinations (43). When fertile eggs are laid, it implies that the embryos in the eggs have already passed the critical point of CI. It has been observed that about one-third of embryogenesis happens within the ovaries before the eggs are laid (24). We expected that if *Rickettsia* induces CI in bed bug, proportion of fertile eggs will be lowest in the group that only males from *Rickettsia*-infected line were crossed (R− x R+).

### Statistical analyses

The data were analysed using the statistical platform R (version 3.6.1, 2019) (44) under the packages ‘lme4’ (45). The analysis of development time and fecundity were done by fitting linear mixed-effects models (LMMs) using the ‘lmer’ function, while sex ratio and cytoplasmic incompatibility were fitted in generalised linear mixed effect model (GLMMs) using ‘glmer’ function with binomial family. For the development time and sex ratio, we fitted ‘pot’ as a random factor of mixed effect models, while *‘*infection status’ and bed bug ‘populations’ were fitted as the main factor in all cases. The proportions of number of female: male and number of fertile: infertile eggs were set as the response variable in the sex ratio and fecundity test, respectively.

In fecundity and CI analyses, ‘infection status’ was broken down into ‘male infection status’ and ‘female infection status’ as these two factors represented cross types (female x male; R+ x R+, R+ x R−, R− x R+, R− x R−), alongside the main factor ‘population’. Family of origin was modelled as a random effect. Non-significant factors were removed from the models until we found the minimum adequate model. We performed Likelihood ratio test (LRT) by comparing the minimum adequate models of both LMMs and GLMMs with a null model using ‘anova’ function, considering by *χ*^*2*^ with critical *p*-value at 0.05. The normality and homoscedasticity of residuals of the LMMs models were validated before the final interpretation.

## RESULTS

### Torix *Rickettsia* across bed bug populations and other cimicids

DNA extractions from each *C. lectularius* population and all cimicid allies passed QC, with good amplicons in the COI amplification. Thirteen out of twenty-one *C. lectularius* populations tested positive to *Rickettsia* infection with both *16S rRNA* and *gltA* primers. These populations have their origin in Africa and Europe (Table 1). Both male and female individuals were found to be infected in most cases where infection was detected. In the initial screen, only the female individua was scored as infected in populations S1 and BG1; however, the infection in males was observed in males in both populations on deeper screening (Table 1). For the cimicid allies, PCR results indicated that only one species, *Afrocimex constrictus* in one subfamily Afrocimicinae, were positive for *Rickettsia* infection with all three samples scoring positive (Table 2). However, the rickettsial PCR only produced *gltA* amplicons for this species, despite repeated attempts at 16S amplification

### Phylogenies

The *16S rRNA* sequences alignment of all *Rickettsia* strains from the *C. lectularius* populations indicated that these strains were identical (based on 985 bp sequence length information, accession number; LR828195). The phylogenetic tree based on *16S rRNA* gene showed the *Rickettsia* strain of *C. lectularius* is placed in torix group (Figure 1). The alignment of *gltA* sequences (746 bp) of *C. lectularius* also showed no variable sites across all *Rickettsia* positive populations. Moreover, the sequences of the *gltA* amplicons from both *C. lectularius* and *A. constrictus* (accession numbers; LR828196-LR828197) were also identical (Figure 1).

**Figure 1.**
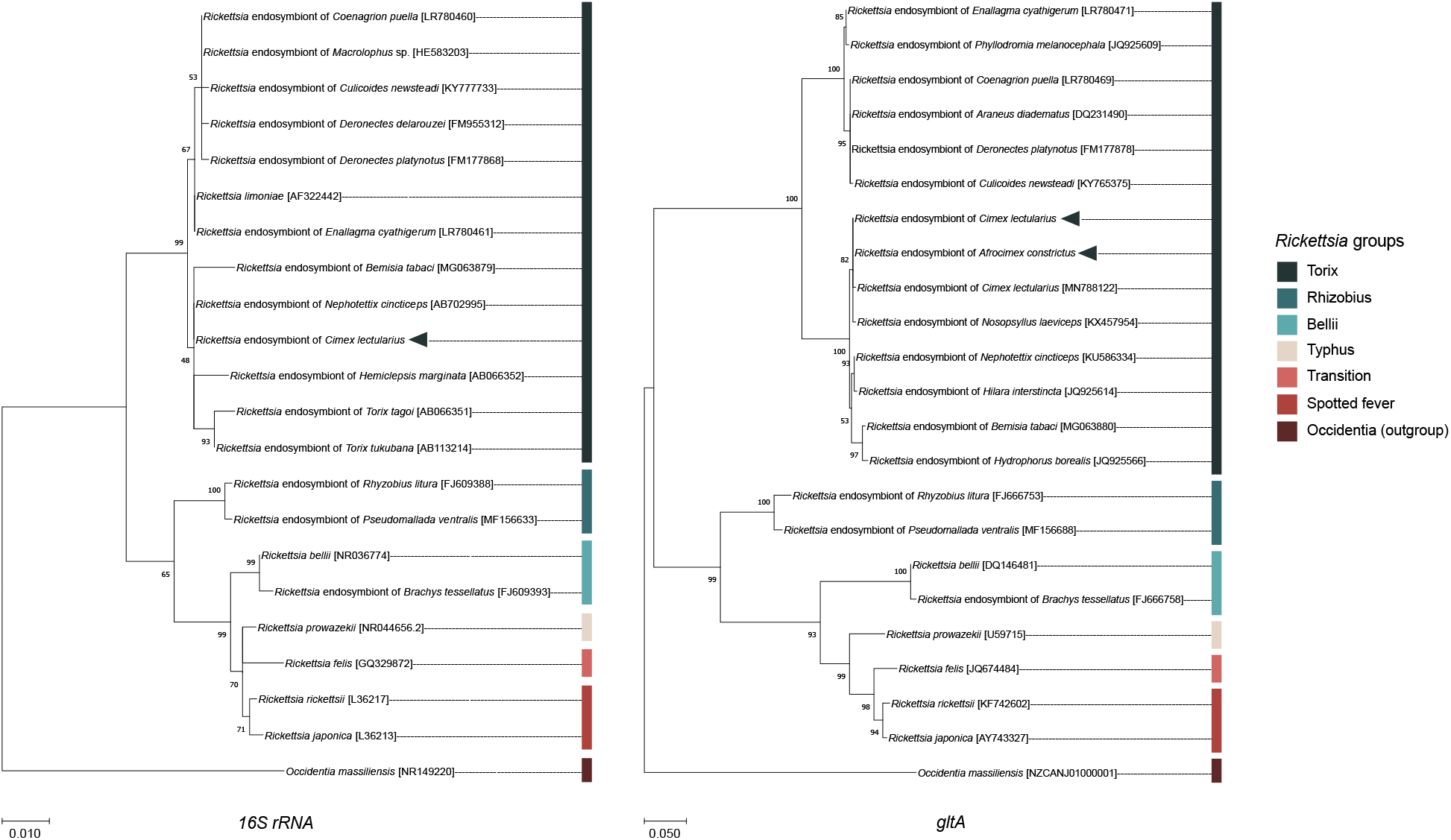
Maximum likelihood phylogenetic trees generated from 16S rRNA gene (A) and gltA gene (B) sequences showing the position and relatedness of *Rickettsia* endosymbiont of *C. lectularius* and *A. constictus* found in this study (arrowhead) with other relative strains that obtained from NCBI (the GenBank accession numbers are in the square brackets). The bootstrap values expressed as the percentage of 1000 replicates are shown at the nodes. Bars indicate substitution nucleotide per position.

### Transmission mode experiment

All 49 crosses of the F4 line produced offspring, while 55 out of 65 crosses from line S1 were successful. The *Rickettsia* infection status for each family was categorised into four groups according to parental infection status. *Rickettsia* transmission to progeny was consistently observed where either both parents (R+ x R+) or just the mother (R+ x R−) tested positive for torix *Rickettsia* (R+ x R+: F4: 8 crosses, S1: 45 crosses; R+ x R−: F4: 6 crosses, S1: 3 crosses) (Figure 2). In these cases, all 310 tested nymphs had acquired infection, indicating vertical transmission through females was highly efficient (Confidence intervals for vertical transmission efficiency 0.988-1.0). No progeny tested positive for *Rickettsia* infection in families where only the father was infected (R− x R+: F4: 12 crosses, S1: 5 crosses), nor were any *Rickettsia* positive individuals recovered from crosses where neither parent was infected (R− x R−: F4: 23 crosses, S1: 2 crosses). These data indicate that maternal infection is essential and sufficient for presence of *Rickettsia* in progeny, and there is no evidence of paternal inheritance. The PCR assay for the BEV-like symbiont revealed that all individuals within these crosses were infected.

**Figure 2.**
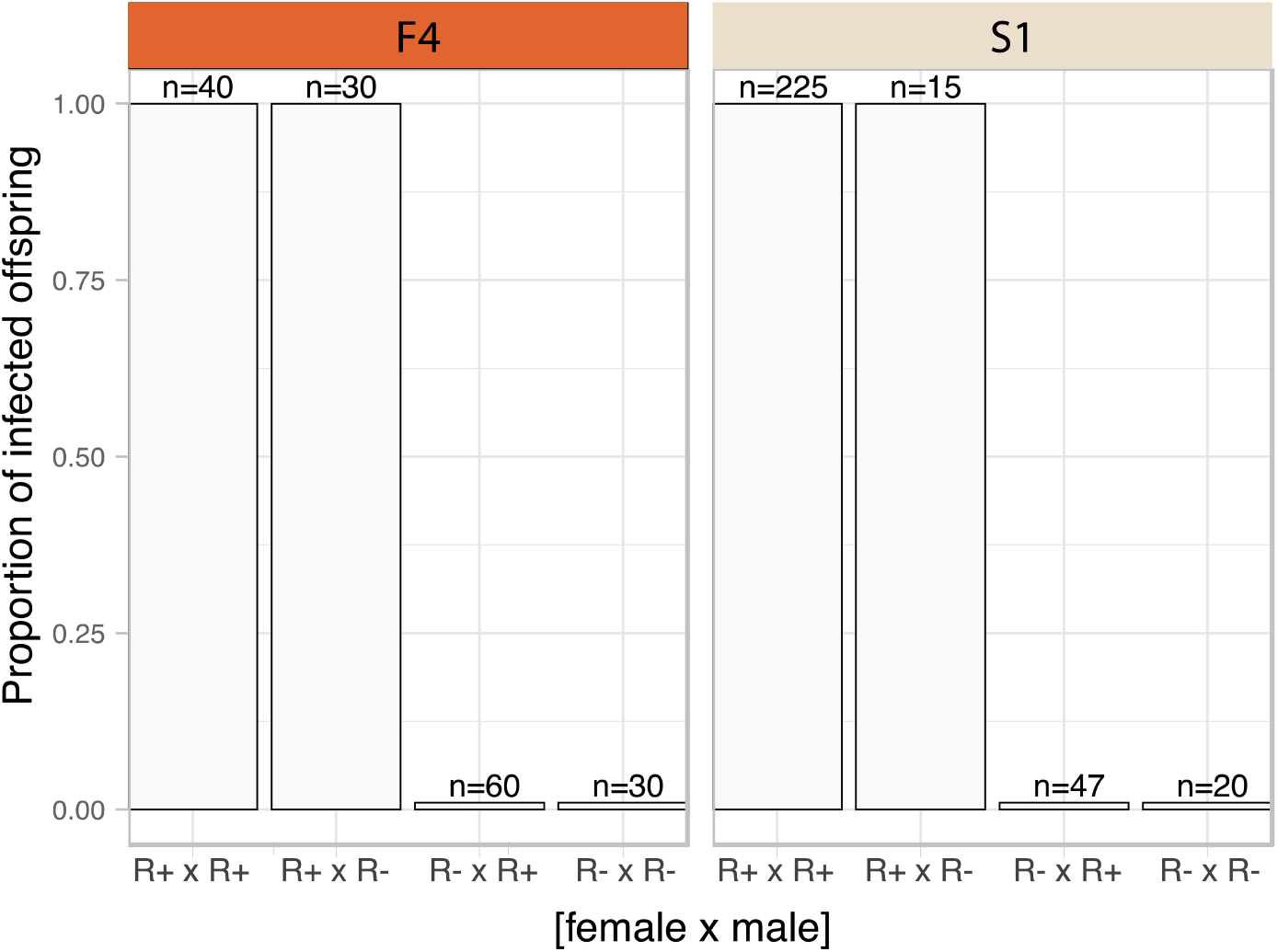
Transmission mode of *C. lectularius* associated *Rickettsia*. The bars indicate proportion of infected offspring in each crossing group. The four crosses were sorted by infection status of the parents, female x male (R+ x R+: 8 crosses for F4, 45 for S1; R+ x R−: 6 crosses for F4, 3 for S1; R− x R+: 12 crosses for F4, 5 for S1; for R− x R−: 23 crosses for F4, 2 for S1). None infected offspring was observed from *Rickettsia*-free mother groups (R− x R+ and R− x R−).

### Localization of the symbionts

FISH detected *Rickettsia* throughout ovaries and bacteriome tissues (Figure 3 and 4 B, D-E). In adult females, the distribution of the *Rickettsia* signal was intense in the trophic core of the tropharium and in oocytes. Alongside *Rickettsia* we found the filamentous BEV-like symbiont and *Wolbachia* in oocytes. In comparison to the other symbionts, *Rickettsia* were more widely distributed in the ovaries. We found them also in nurse cells and follicular epithelial cells (Figure 3 C-D). *Rickettsia* signals were absent in *Rickettsia*-free bed bugs (Figure 3 E-F).

**Figure 3.**
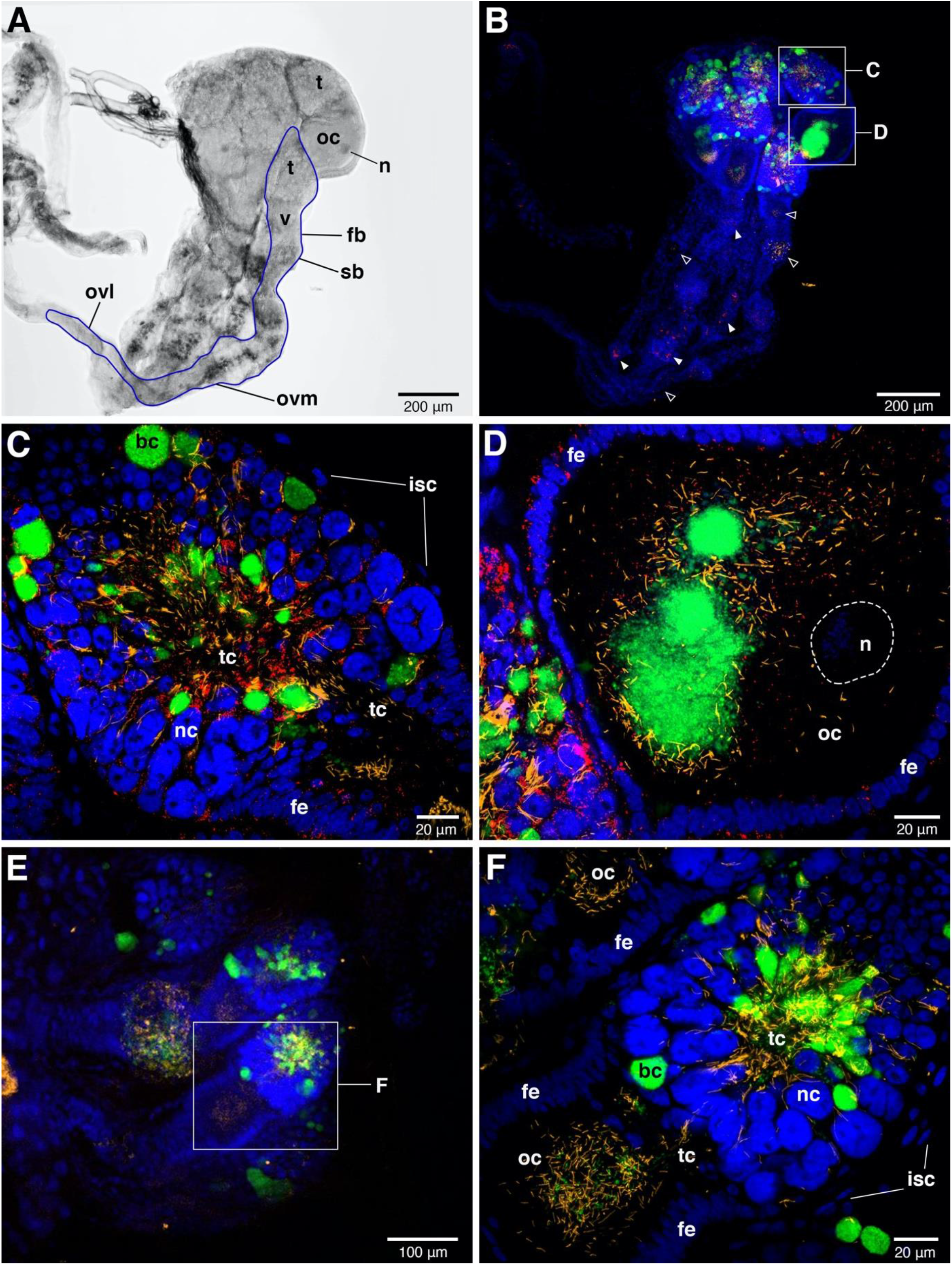
FISH images of adult female ovaries. **A:** Bright field image of one-sided ovaries from torix-infected female. The blue line represents the outline of one ovariole. **B**: FISH shows the present of the three symbionts i.e. *Wolbachia* (green), BLS (yellow) and *Rickettsi*a (red) in ovaries. These signals are concentrated in tropharium areas. Small rectangles indicate the magnified fields of tropharium and vitellarium portions that are showing in **C** and **D**, respectively. *Rickettsia* and BLS are also detected in syncytial body and mesodermal oviduct at low densities, the present of the signals are shown in filled and clear arrowheads, respectively. **C**: The enlarge detail of tropharium portion. The three symbionts distribution can be detected at very high density in all along the trophic core area. *Wolbachia* (green) are likely packed in bacteriocytes (bc) which are distinctive to the adjacent nurse cells (nc), while *Rickettsia* and filamentous BLS are more scattered. **D**: The enlarge detail of vitellarium portion. All the three symbionts invade in oocyte, forming a cluster at the presumable posterior pole of the oocyte. *Rickettsia* signals are scattered insertion in the follicular epithelium (fe) of the oocyte. **E**: Ovaries of torix-free bed bug. Only *Wolbachia* and BLS are present. The small rectangle represents the tropharium and vitellarium parts of one ovariole showing in **F. F**: The enlarged field of the upper ovariole partial. The distribution of *Wolbachia* and BLS are intensive in trophic core and in oocyte. None *Rickettsia* signals are detected here. bc = bacteriocyte, fe = follicular epithelium, fb = follicular body, isc = inner sheath cell, n = nucleus of oocyte, nc = nurse cells, oc = oocyte, ovl = lateral oviduct, ovm = mesodermal oviduct, sb = syncytial body, t = tropharium part, tc = trophic core, v = vitellarium part.

**Figure 4.**
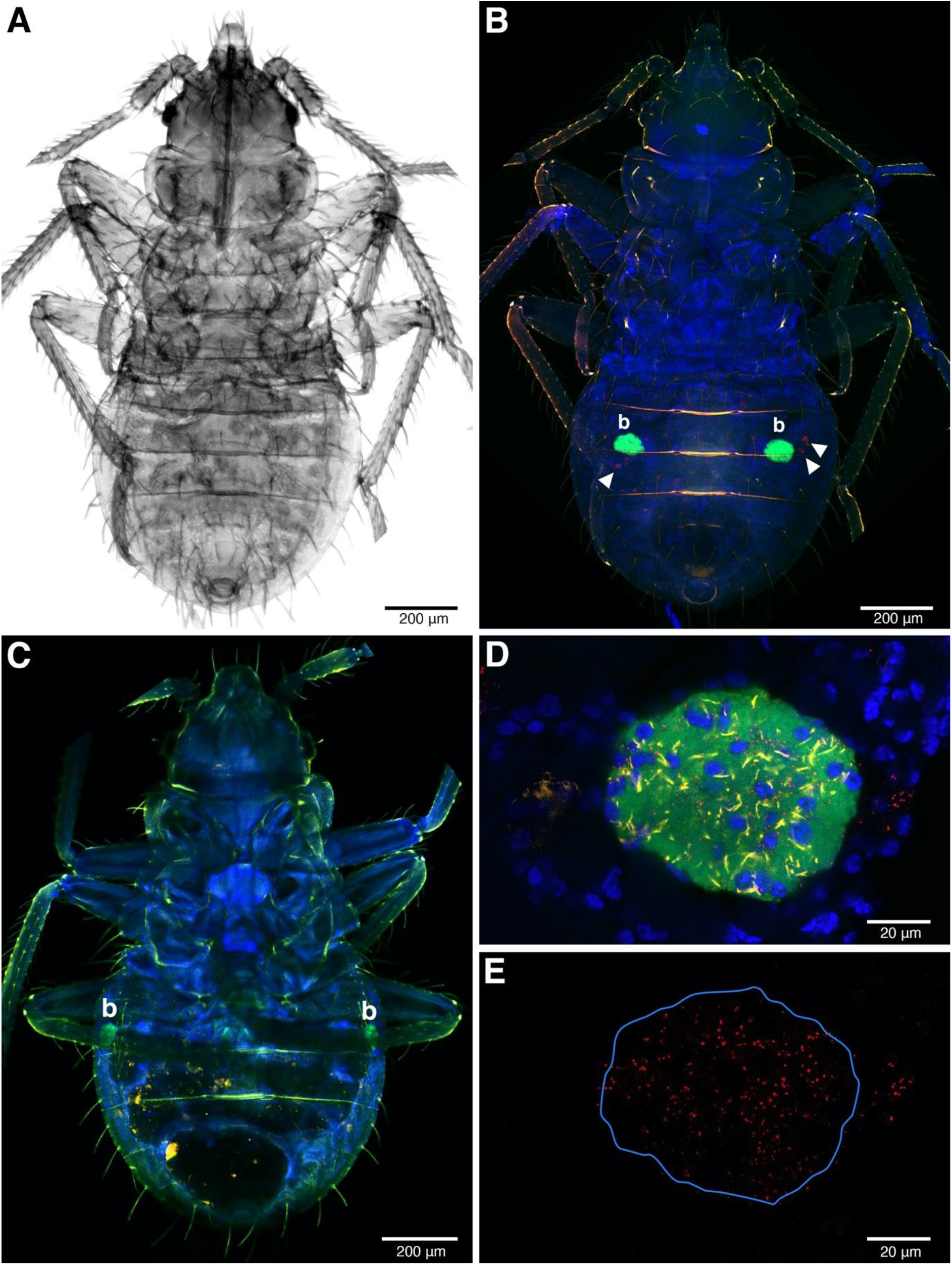
FISH image of the whole-mounted first instars. **A**: The nymph under transmitted light. **B-C**: FISH detection of symbionts in torix-infected (**B**) and torix-free instars (**C**). The ball-shaped in green colour represents strong *Wolbachia* signals indicate where bacteriome allocation in abdomen, *b* = bacteriome. The *Rickettsia* infection showing in red where the white arrowhead present in **B** but absent in **C. D**: Bacteriome of torix-infected instar. All the three symbionts can be detected in this organ. *Wolbachia*, BEV-like symbionts and *Rickettsia* are in Green, yellow and red respectively. The blue colour represents nuclei of bed bug cells. **E**: The same field as **D** but only leave *Rickettsia* channel remains. The blue line indicates the bacteriome boundary.

*Rickettsia* signals were found in somatic tissue of the abdominal areas of first instar whole mounts, alongside the BEV-like symbiont (Figure 4 B). At low magnification, the strongest signal (green colour) was emitted by *Wolbachia* reflecting the intense infection of bacteriome organs (Figure 4 B). *Rickettsia* and the BEV-like symbiont, however, could also be seen at this low magnification but poorly resolved. At the higher magnification, *Rickettsia* is clearly visible in the bacteriome alongside *Wolbachia* and the BEV-like symbiont as all the three signals were reliably present in this tissue (Figure 4 D-E). In *Rickettsia*-free samples only the signals of *Wolbachia* and the BEV-like symbiont were present, while *Rickettsia* signals were absent (Figure 4 C).

### Development time and sex ratio

The development of *C. lectularius* was slowed by infection with torix *Rickettsia* (LRT: *χ2*(1) = 5.177, *p* = 0.023) with first instar-adult period for *Rickettsia* infected individuals increased by 0.59±0.26 days (Mean±SD). The *Rickettsia*-infected line took 26.7±2.00 days to reach to adulthood (N = 78) while the bugs from the *Rickettsia*-free line took 25.9±2.15 days (N = 72) (Figure 5). Comparison between the model with interaction and non-interaction terms of the fixed factors (infection status, sex and populations), likelihood ratio indicated that there were no interaction effects between those factors on development time of the *C. lectularius* (LRT: *χ2*(3) = 5.232, *p* = 0.156).

**Figure 5.**
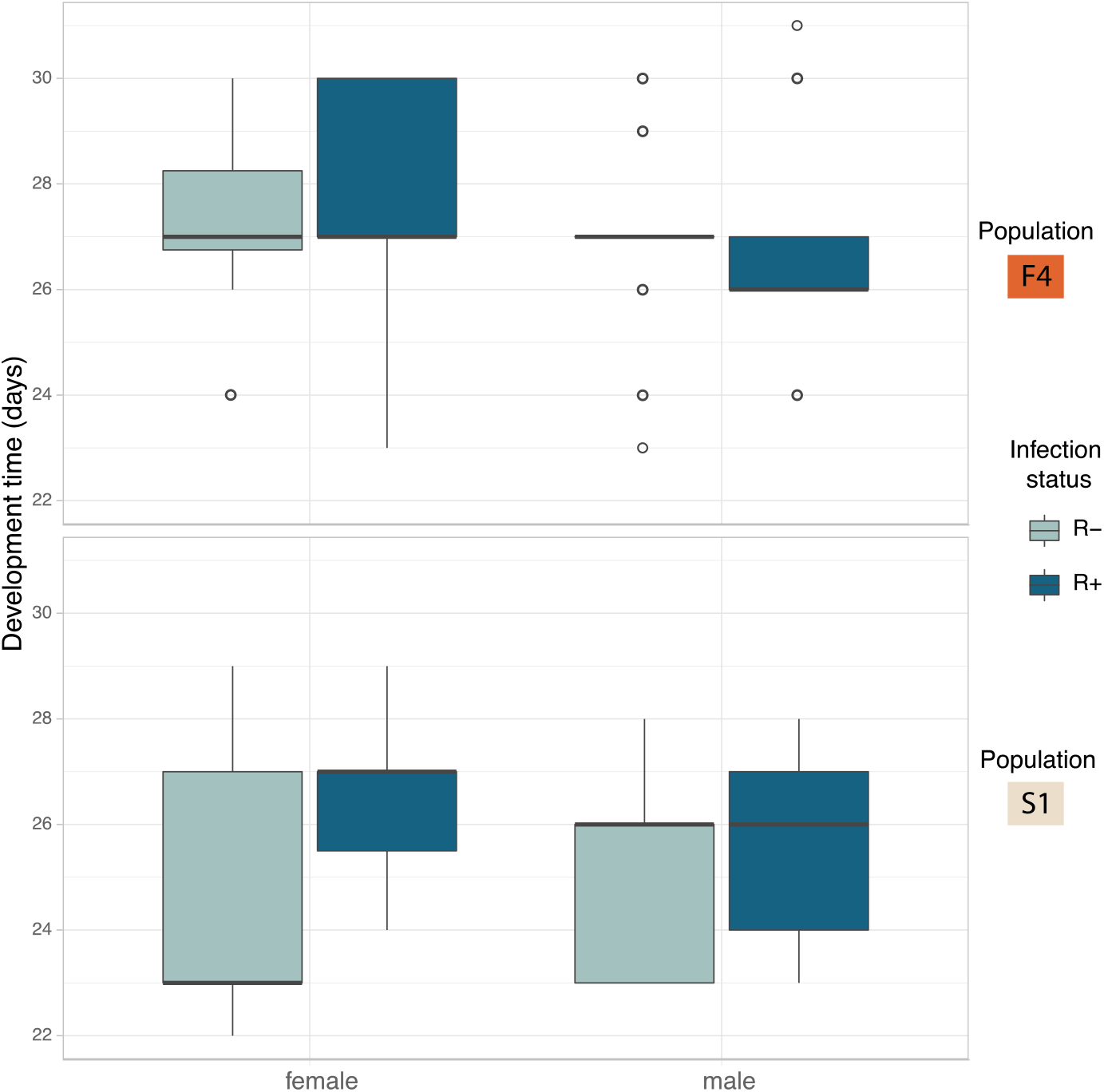
Development time in days of *C. lectularius* from the first instar to adulthood for males and female individuals of *Rickettsia*-free (R−) and *Rickettsia*-infected (R+) groups from population F4 and S1. *Rickettsia* infection has a significant effect on development time (LRT: *χ2*(1) = 5.177, *p* = 0.023).

There was no impact of *Rickettsia* infection status on sex ratio of offspring (LRT: *χ2*(1) = 0.0003, *p* = 0.985) (Figure 6). Female: male ratio of the *Rickettsia*-free group was 0.84±0.29 and 0.93 ±0.67 for the *Rickettsia*-infected group. There was no interaction effect between infection status and population of origin (LRT: *χ2*(1) = 0.078, *p* = 0.780).

**Figure 6:**
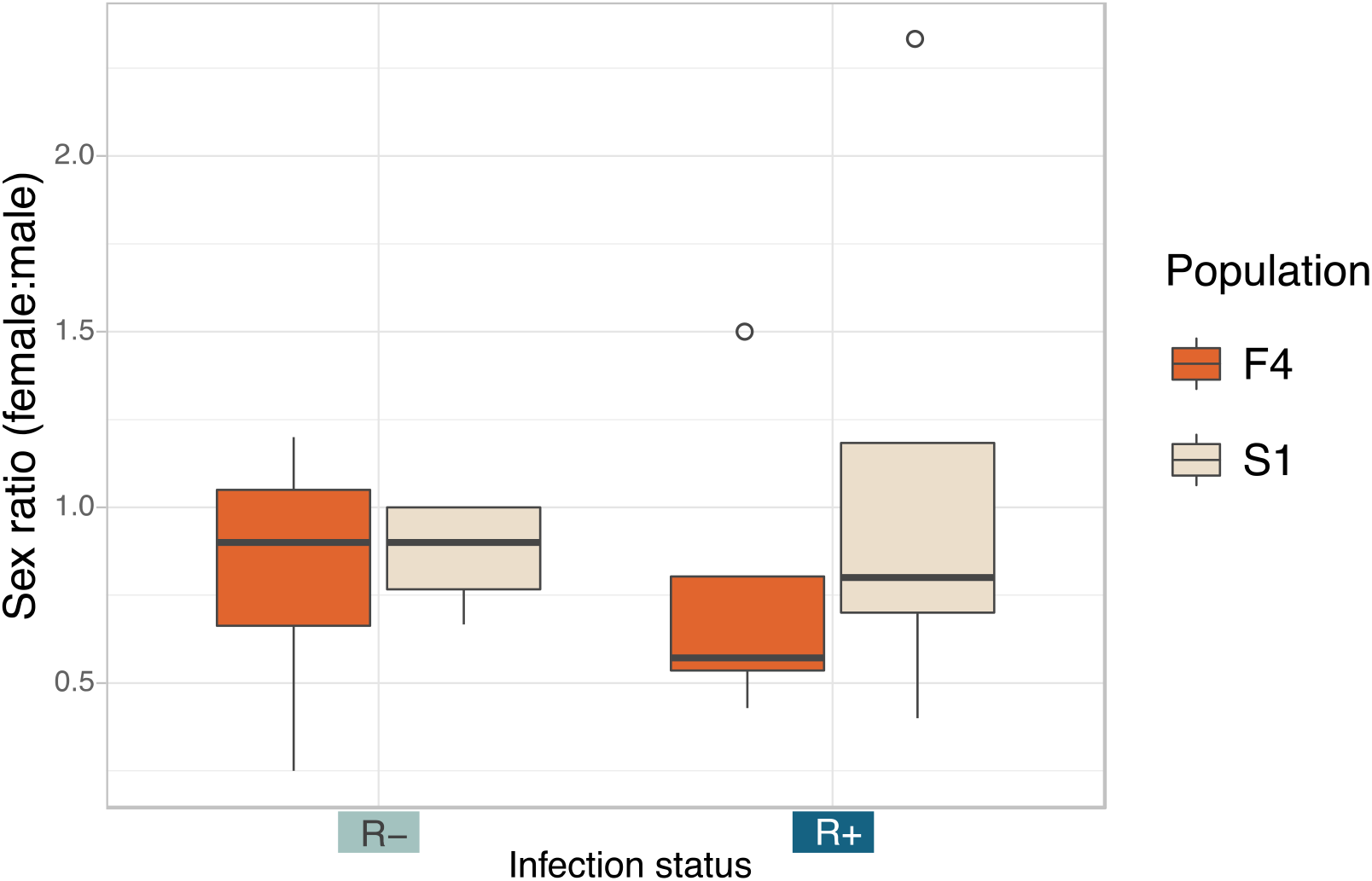
Sex ratio (number of female:male) of the *Rickettsia*-free and *Rickettsia*-infected *C. lectularius* adults from the two populations. The sexes were identified from adult bed bugs. There was no significant different of the sex ratio between R− and R+ group at *p* = 0.05.

### Fecundity and Cytoplasmic incompatibility

We analysed fecundity (total fertile eggs) using Likelihood ratio comparison of LMMs. There was no evidence of an interaction effect between the three factors, i.e., population, male and female infection status (LRT: *χ2*(3) = 5.781, *p* = 0.216), and these terms were dropped from the model. The final model detected *Rickettsia* infection in female parent as the sole significant explanatory variable for fecundity (LRT: *χ2*(1) = 4.576, *p* = 0.032). Infected females were likely to produce fewer fertile eggs (R+ x R+ = 86.80±34.30, R+ x R−= 89.70±28.30) compared to non-infected females (R− x R+ = 107.00±34.90, R− x R− 109.00±34.60, Figure 7 A).

**Figure 7:**
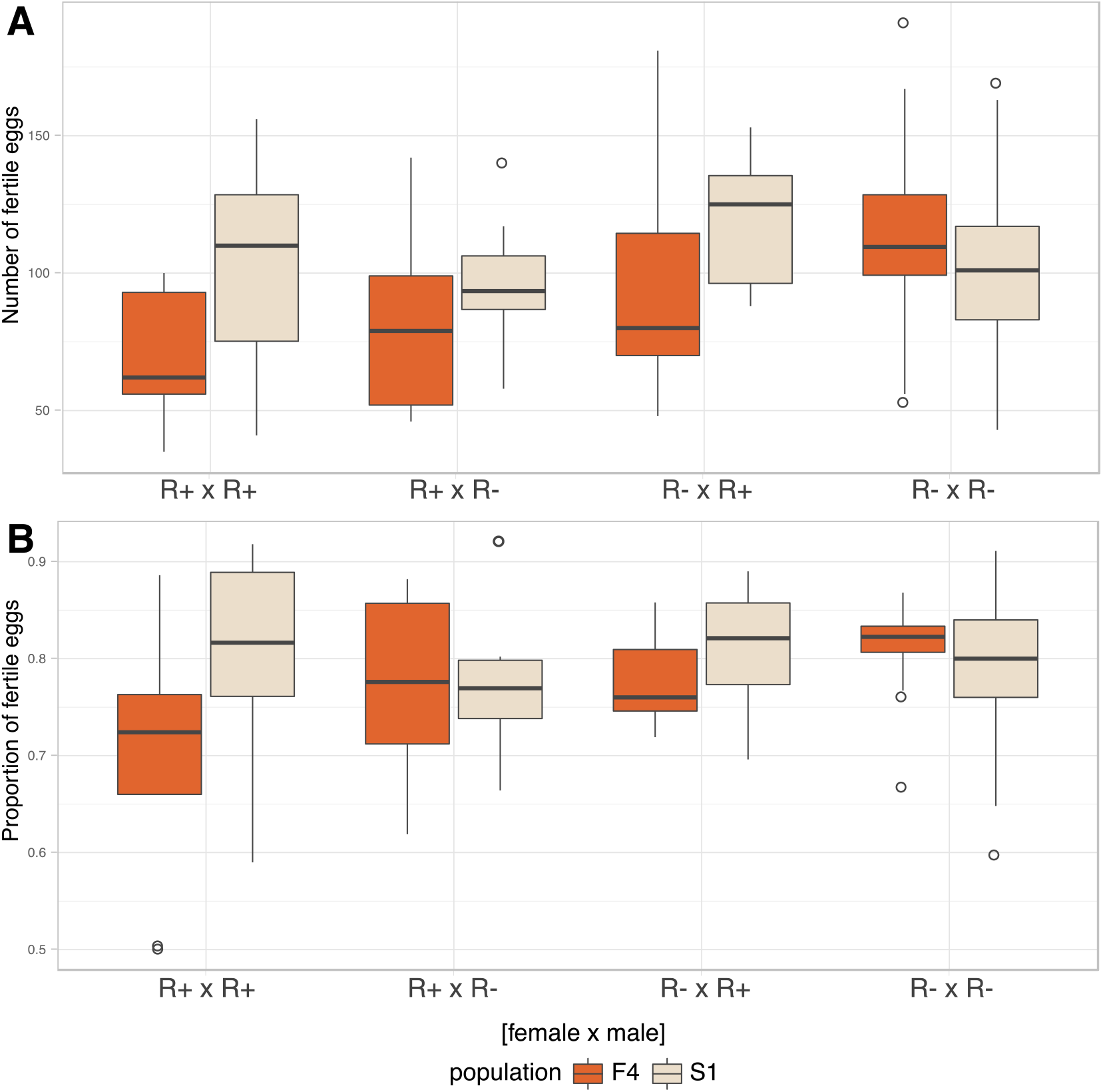
Fecundity and CI. **A**: Number of fertile eggs of *C. lectularius* from population F4 and S1 from the four cross combinations. *Rickettsia*-infected female crosses (R+ x R+ and R+ x R−) produced significantly fewer fertile eggs when compare to other crosses (LRT: *χ2*(1) = 4.5761, *p* = 0.032). **B**: Proportion of fertile eggs of *C. lectularius* from the two populations. There was no interaction effect of male and female infections on the ratio of fertile:infertile eggs from the GLMMs analysis at *p* = 0.05, indicating there was no evidence of CI.

We then analysed the relative ratio of fertile:infertile eggs to ascertain if there was any evidence of cytoplasmic incompatibility. There was no evidence of heterogeneity associated with *Rickettsia* infection in either male (LRT: *χ2*(1) = 0.593, *p* = 0.441) or female parents (LRT: *χ2*(1) = 1.174, *p* = 0.279). There was no evidence of an interaction term between male infection x female, evidence by the statistical equivalence of models with and without an interaction term (LRT: *χ2*(4) = 6.725, *p* = 0.151, Figure 7 B).

## DISCUSSION

Torix *Rickettsia* are common associates of insects and other invertebrates, but their mode of transmission and impact on the host are poorly understood. Intriguingly, in the one case where transmission modes has been explicitly examined, transmission through both male and female lines was observed (46). In the present study, we examined the interaction between torix *Rickettsia* and *C. lectularius*, a notorious pest of humans. In line with other recent work, our PCR screen revealed *Rickettsia* were commonly found in bed bug lines maintained in the laboratory. In our study, the observation revealed only a single *Rickettsia* strain. The segregation of the *Rickettsia* in two lines was observed – with a mix of infected and uninfected individuals being observed in laboratory populations. The polymorphism in *Rickettsia* presence allowed us to isolate isogenic R+ and R− cultures and then use these to analyse transmission and impact on the host. We observed that the symbiont showed high fidelity maternal inheritance, but no transmission through males. Consistent with this, *Rickettsia* infections were present in ovaries, as well as in the bacteriome, and more diffusely. Infection with this symbiont did not impact host sex ratio and did not induce cytoplasmic incompatibility, but there was evidence it slowed development to the adult stage and reduced female fecundity.

In this study, the *Rickettsia* strain detected in *C. lectularius* and *A. constrictus* are placed in the non-vertebrate pathogen ‘torix group’ which consisted of *Rickettsia* endosymbionts of other arthropods, e.g., *Nosopsyllus* flea, *Culicoides* biting midge and other non-arthropod hosts e.g. glossiphoniid leeches (*Hemiceplis marginata, Torix tugubana* and *T. tagoi)*. The detection of a single torix *Rickettsia* strain in cosmopolitan bed bug species indicates that one main strain of this symbiont circulates in this species worldwide, consistent with movement alongside travellers. This finding also appears in the recent study of Potts et al., (28). These *Rickettsia* strains were investigated in *C. lectularius* from UK, which are independent from the populations in our study, and field collections from the USA, all of which reveal an identical *gltA* haplotype.

It is typical for inherited endosymbionts to be transmitted maternally between host generations (47, 48). Similarly, our bed bug associated torix *Rickettsia* was observed to be maternally inherited. In the present experiments, maternal inheritance reliably occurred in 100% of the tested offspring. However, the segregation of the symbiont during 12 years of laboratory passage (ca. 100 bug generations) in line F4 indicate some level of inefficient maternal inheritance. Thus, we can conclude that whilst vertical transmission through females is very high, it does not occur with a 100% efficiency.

In contrast, no paternal transmission was observed in this study, despite paternal males carrying infections, and previous evidence of *Rickettsia* in male sperm vesicles (49). The situation contrasts with the leafhopper-associated *Rickettsia*. In *Nephotettix cincticeps*, torix *Rickettsia* can transmit biparentally, with 70% paternal and 100% maternal transmission rates. The *Rickettsia* are found in sperm without interrupting sperm function (46). Paternal inheritance has also been noted for symbionts in the genus *Megaira*, the sister taxa to *Rickettsia* (50). Notably, the *Rickettsia* in the leafhopper has the capacity for intranuclear infection, which likely is necessary for paternal inheritance. Intranuclear infection is present quite widely in the Rickettsiacae, but is labile, being present in some symbioses but not others (51).

Insects commonly live mutualistically with endosymbionts to facilitate each other. In many cases, insect hosts provide a pair of bacteriomes as organs for harbouring their endosymbionts (3, 48, 52-54). In previous histological studies, the bacteriome organs are located at either side of the bed bug abdomen. So far, only two bacteriome-associated endosymbionts have been described in *C. lectularius*, i.e. the primary endosymbiont *Wolbachia* and a secondary BEV-like symbiont (3). Here we additionally examined the tropism of *Rickettsia*, alongside *Wolbachia* and the BEV-like symbiont in different organs. As expected *Wolbachia* dominated the bacteriome in terms of numbers. *Rickettsia* was also present in the bacteriome and the three symbionts were spatially intermixed within them. This result contrasts with the localization the symbiont community in the leafhopper *N. cincticeps*, in which its symbionts live separately. The facultative *Rickettsia* is widely dispersed through *N. cincticeps* bacteriomes and can be found in most of the leafhopper tissues (55). When the symbiont infection has a specific territory within the bacteriome this implies a more specific function of those symbionts for their host (56, 57). Investigating in-depth in the bed bug bacteriomes might help to understand the distribution of these three endosymbionts and could potentially anticipate the biological impacts of these endosymbionts on the host or their interaction.

The presence of *Rickettsia* in bacteriocytes indicates a route for achieving maternal inheritance. In this system vertical transmission of *Wolbachia* is associated with the movement of bacteriocytes towards the ovary, where they fuse with oocytes to deliver the symbiont. The presence of *Rickettsia* in the bacteriome likely allows this symbiont to hitch-hike to the ovary to gain vertical transmission. However, having established in the ovary, *Rickettsia* and *Wolbachia* show distinct patterning, with *Wolbachia* clustering in discrete clumps and *Rickettsia* being more dispersed.

We also examined the impact of *Rickettsia* on bed bug biology. We found no evidence of reproductive parasitic phenotypes (sex ratio distortion or cytoplasmic incompatibility). There was evidence of a weak negative impact on first instar-adult development time, and also reduced fecundity of *Rickettsia* infected females. Overall, all metrics that we examined showed either no evidence of presence, or a deleterious impact of *Rickettsia* on the property.

The combination of segregational loss and modest costs indicate maintenance of the symbiont in field populations will require some balancing benefit. A recent example of the biological impact of torix *Rickettsia* can be found in glossiphoniid leeches-associated *Rickettsia*. The case study demonstrates that *Rickettsia* have a direct effect on the body size of the three leech host species, with infected individuals being larger (16). A recent study has examined the genome sequence of torix *Rickettsia* endosymbiont of *Culicoides newsteadi*, a strain closely related to bed bug-associated *Rickettsia* (17). No evidence of the capacity for positive facilitation was observed in the genome, e.g. B-vitamin provisioning. However, the genome possesses unique features of genes that are potentially associated with host invasion and adaptation (17). It is likely that any benefits of *Rickettsia* infection are ecologically contingent, and parallels with other symbioses indicates resistance to environmental stress or pathogen susceptibility as worthwhile avenues for research.

## CONCLUSIONS

This study confirmed the existence of a *Rickettsia* associate in a blood-feeding insect, a potential disease vector. The strain identified is in the ‘torix group’, in which none of the members are known to be vertebrate pathogens. The result revealed the consistent of single *Rickettsia* strain circulates in the cosmopolitan *C. lectularius* populations and this strain is also identical to the *Rickettsia* in an African bat bug (*A. contrictus*). Maternal inheritance was demonstrated, but there was no evidence of reproductive manipulation phenotypes (CI, sex ratio distortion). There was evidence that *Rickettsia* infected individuals had modestly delayed development and that *Rickettsia* infected females laid fewer fertile eggs. The combination of segregational loss, weak physiological cost and absence of reproductive parasitism suggests the presence of a balancing, but yet to be discovered, beneficial effect. Further studies on how this symbiont alter host benefits are worth to explore as well as other dimensions of the symbiosis biology still remain for investigations.

## ACKNOWLEDGEMENTS

We are grateful to Steffen Roth (University Museum of Bergen) and Klaus Reinhardt (TU Dresden) who made samples available from their recently published phylogeny (Roth et al. 2019). The project was supported by the Development and Promotion of Science and Technology Talents Project (DPST) of the Institute for the Promotion of Teaching Science and Technology, Thailand to PT. OO was supported by a grant from the German Research Foundation DFG (OT 521/2-1). Microscopy was conducted at the Centre for Cell Imaging (CCI), using equipment funded from BBSRC grant BB/M012441/1.

## Data deposition

Sequences have been deposited in EMBL (Accession numbers; LR828195-LR828197). Data underpinning Figures 5-7 can be accessed at https://doi.org/10.6084/m9.figshare.c.5127290.v1

---

**Supplementary Table 1.**
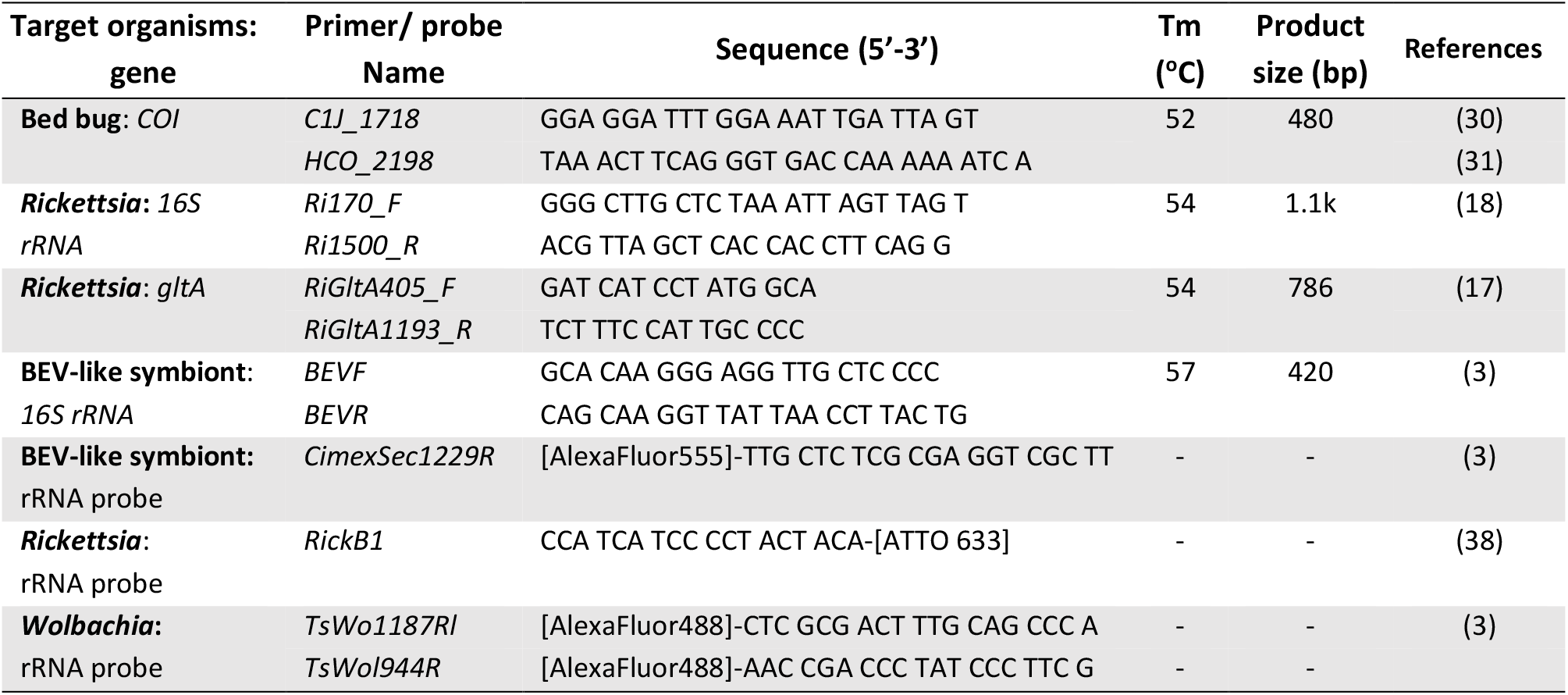
PCR primer and fluorescence probe sequences that were used in this study. All the fluorescence probes are labelled with fluorophore, in the square brackets, at 5’ end except *RickB1* probe, labelled at 3’ end. All the primers were used in the following PCR conditions; initial denaturation at 95 °C for 5 min, followed by 35 cycles of denaturation (94°C for 30s), annealing (Tm°C for 30s), extension (72°C for 50s), and a final extension at 72°C for 7 min. The annealing temperature was varied according to the primers.

